# Identification of the Goliath beetles *Fornasinius fornasini* Bertoloni and *Fornasinius hauseri* Kraatz (Scarabaeidae: Cetoniinae)

**DOI:** 10.1101/242750

**Authors:** Michele De Palma, Hitoshi Takano

## Abstract

The Goliath beetles (subtribe *Goliathina*) are large Cetoniinae comprising a few highly related genera broadly distributed in sub-Saharan Africa. The genus *Fornasinius* Bertoloni was established in 1853 by Giuseppe Bertoloni to receive a taxon that was sufficiently distinct from the known representatives of the genus *Goliathus* Lamarck to merit a different placement. In spite of their large size and showy appearance, the members of the genus *Fornasinius* are poorly known. Here, the type species of the genus, *F. fornasini* Bertoloni, 1853 (type locality: Mozambique), is identified and re-described. It has emerged that *F. fornasini* Bertoloni has been misidentified after its original description and that only a few specimens are known of this species. *F. fornasini sensu auct*. (*nec* Bertoloni) is instead referable to another taxon, *F. hauseri* Kraatz, 1896 *sp. bon.*, of which three subspecies can be distinguished: *F. hauseri s. str.*, from south-central Kenya and possibly Cameroon; *F. hauseri* ssp. *hirthi* Preiss, 1904 *stat. rev.*, from the Lake Victoria region and northern Tanzania; and a new subspecies to be described from south-east Democratic Republic of the Congo.

## Results and discussion

### Genus *Fornasinius* Bertoloni, 1853

#### Type species: *Goliathus fornasini* Bertoloni, 1853

Bertoloni (1853) proposed the generic name ***Fornasinius*** to receive a new species of *“Goliathus”* collected in Inhambane, southern Mozambique (*“Habitat in palmetis riparum fluminis Magnárra Mozambici”*). The description of *Goliathus Fornasini* Bertoloni was based on four specimens of both sexes (*“Di questo animale mi furono mandati quattro individui perfettissmi, uno maschio, e tre femine*”; Bertoloni, 1853), two of which, a male and a female, have been illustrated (Fig. 1).

**Fig. 1.**
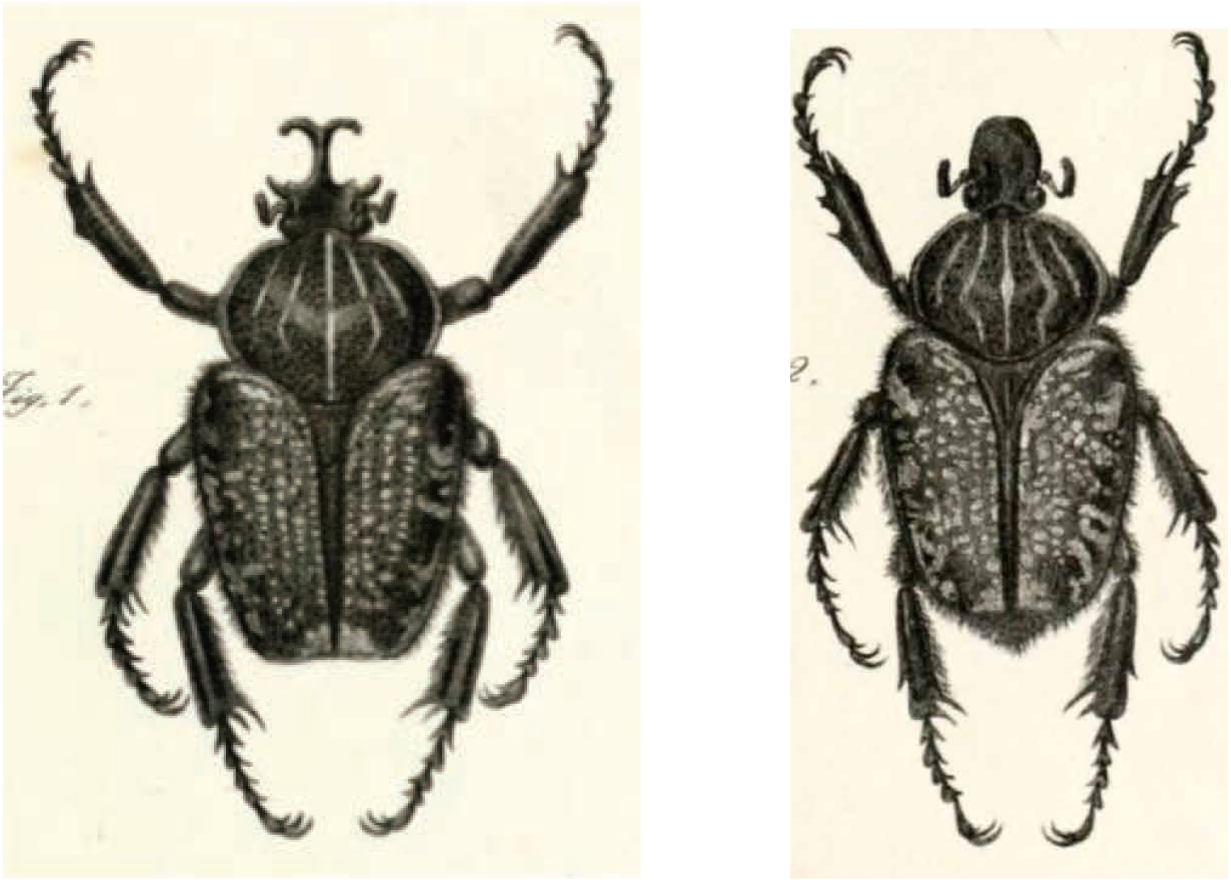
Original illustrations of the only male syntype (left) and a female syntype (right) of *Fornasinius fornasini* Bertoloni (Bertoloni, 1853).

In his paper, Bertoloni noted the proximity of the new taxon to the three *Goliathus* species known at the time, which in his view justified the cautionary placement of the new taxon in Lamarck’s genus *Goliathus*. However, he also noted differences that could justify the creation of a new genus. Bertoloni recommended *Goliathus fornasinius* be renamed ***Fornasinius insignis***, should the creation of a new genus be envisioned by subsequent authors (“*mentre in essa la maggior parte dei caratteri generici dedotti da ambi i sessi sono di Goliathus, il maschio ne presenta alcuni altri di tale importanza e non comuni colle tre specie conosciute summenzionate, che forse, come dissi di sopra, si potrebbe innalzare al rango di genere novello, vicinissimo al Goliathus.”* […] “se *mai si dovesse ricevere in un genere nuovo, io bramerei, che ad onore del ritrovatore benemerito fosse dagli entomologi distinto col nome di Fornasinius insignis; cosi la scienza si mostrerebbe grata alle premure di chi la fa progredire.”;* Bertoloni, 1853). In doing so, Bertoloni explicitly established a new genus name, *Fornasinius*, with the type species *Goliathus fornasini*. The specific name *Fornasinius insignis* must, therefore, be considered a synonym of *Fornasinius fornasini*, although some authors (e.g., Kolbe, 1913) treated the new taxon as *Fornasinius insignis*.

Thomson (1856) re-described *Goliathus Fornassinii* (misspelling of *fornasini*) and illustrated a female specimen (see below), without adopting the new generic name proposed by Bertoloni (*“Cette espèce doit être placée auprès des G. giganteus, Druryi et cacicus, peut-être après le G. Druryi”;* Thomson, 1856).

Westwood created the name ***Goliathinus*** as a subgenus of *Goliathus* to receive *Goliathus fornasini* (Westwood, 1874), thus neglecting Bertoloni’s recommendation to use *Fornasinius* as a new generic name. Therefore, *Goliathinus* is an objective synonym of *Fornasinius*. In his description, Westwood cites Bertoloni’s type as well as a male specimen from the Turner collection, which he illustrated (see below). Subsequent authors, with few exceptions, treated Bertoloni’s taxon as *Fornasinius fornasini*, which is the correct combination for this species.

### Identification of *Fornasinius fornasini* Bertoloni, 1853

The taxon currently known as *Fornasinius fornasini sensu auct.*, which is distributed in the surroundings of Lake Victoria and north-east Tanzania, is not consistent with Bertoloni’s description and illustrations of *Fornasinius fornasini;* its identity will be clarified below. Two syntypes of *Fornasinius fornasini* could be located in the Dipartimento di Scienze Biologiche, Geologiche e Ambientali of the Universitá di Bologna (UNIBO), Italy. Prof. Mario Marini kindly communicated photos of the syntypes (a male and a female), which unambiguously correspond to the two syntypes illustrated by Bertoloni.

While the habitus of the *Fornasinius fornasini* syntypes is reminiscent of that of *Fornasinius fornasini auct*. (*nec* Bertoloni), the cephalic armature of the male shows striking differences (Fig. 2). The horn of *Fornasinius fornasini* erects from the frons and projects antero-dorsally, without curving ventrally as in *Fornasinius fornasini auct*. (*nec* Bertoloni). It is compressed laterally along its entire length; its base is not as dilated as in the other members of this genus; and its terminal fork has upturned spines. Bertoloni provided a detailed description of the cephalic horn of *Fornasinius fornasini: “La forma sua è singolare, perché dalla sua allargata base alzasi gradatamente restringendosi, e mostrasi compressa ai lati per tutta te sua lunghezza col bordo posterior convesso, coll'anteriore protuberante in un angolo coll'apice rotondato ed i lati quasi retti, angolo collocato alla metà della lunghezza dello stesso bordo. L'apice suo poi si biforca in due cornetti rotondati, coll'estremità ottusa, allungati e divaricati verso i lati della testa, superiormente divisi l’uno dall'altro da un profondo e largo solco.”* (Bertoloni, 1853). An obtuse cornicle projecting anteriorly is present at each side of the horn.

**Fig. 2.**
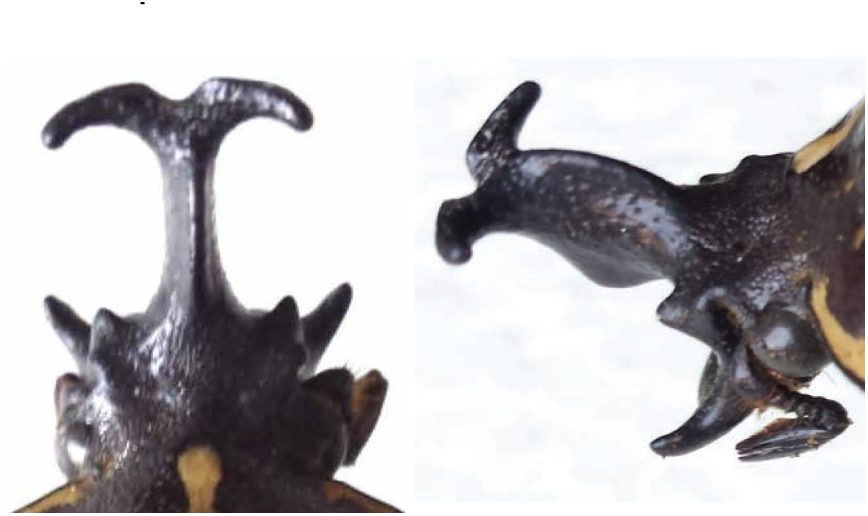
Head of the male syntype of *Fornasinius fornasini* Bertoloni, housed in UNIBO.

The clypeal plate of the head (Fig. 3) is similar to that of *Fornasinius fornasini auct*. (*nec* Bertoloni); each side of it is armed with an elongate, strongly indented spine, whose tip is truncated transversally. The lateral spines of the clypeal plate are clearly visible from above (see Fig. 2).

**Fig. 3.**
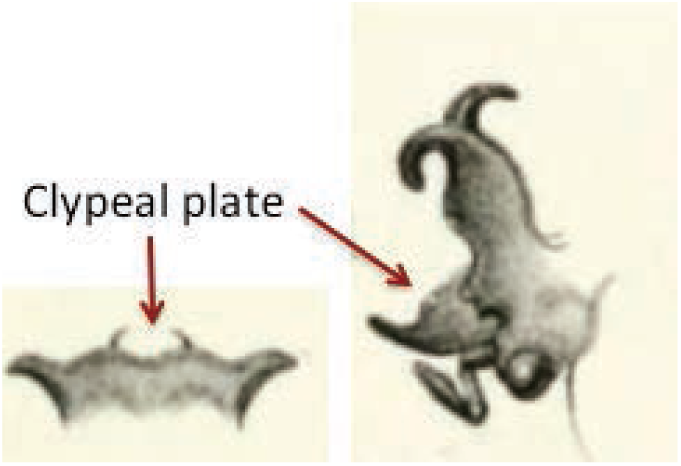
Annotated original illustrations of the head of the male syntype of *Fornasinius fornasini* Bertoloni (Bertoloni, 1853).

Conversely, the horn of *Fornasinius fornasini auct*. (*nec* Bertoloni) has a broad, triangular base; in large specimens, it erects almost vertically from the frons to abruptly deviate ventrally at about a third of its length. Diminutive specimens of *Fornasinius fornasini auct*. (*nec* Bertoloni) may display short and straight horns, but this structure is never observed in specimens of medium to large size. Notably, the male syntype of *Fornasinius fornasini* is a very large specimen, measuring 58 mm from the tip of the horn to the tip of the elytra.

In addition to the two certain syntypes (the remaining two female syntypes mentioned by Bertoloni are not in UNIBO), only a few other specimens consistent with Bertoloni's taxon could be found.

A female specimen collected in Mozambique was illustrated by Thomson (1856) under the name of *Goliathus Fornassinii* (misspelling of *fornasini*). This specimen, which carries a type label written in Thomson’s hand (“*Insignis* / Bertol. Typ. D / *Fornassinii* An. 1856 / Bert. *Olim*. / Mozamb.”), was located in the Muséum national d'Histoire naturelle (MNHN) of Paris, France (Fig. 4). Thomson (1856) wrote that he had been entrusted with the specimen by Dohrn (*“qui m'a confié cet insecte jusqu'à son retour d'Italie”*), and that two additional female specimens were present in his own collection (not traced). It is possible that Dohrn’s specimen had belonged to Bertoloni’s type series.

**Fig. 4.**
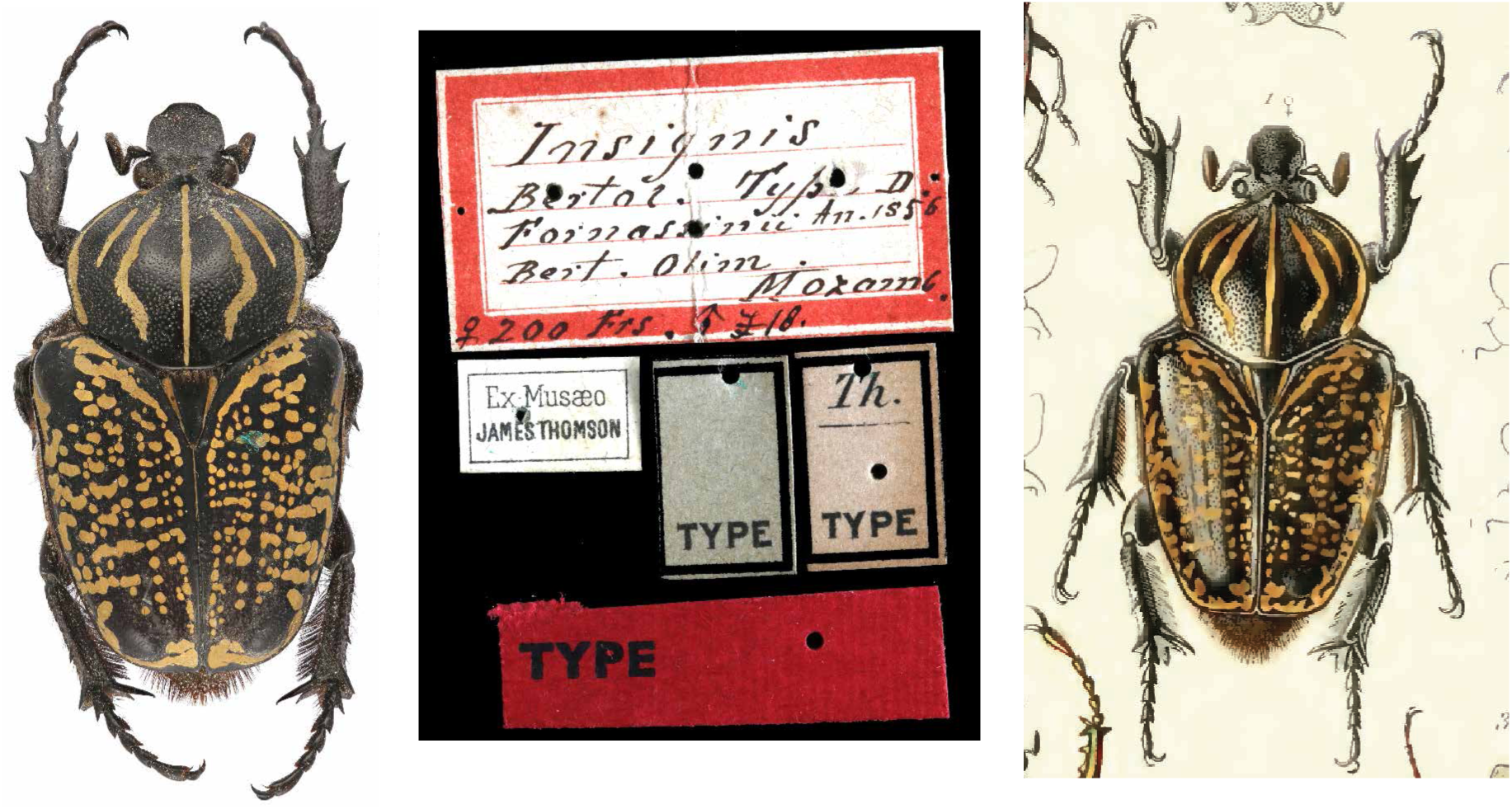
Thomson's specimen of *Fornasinius fornasini* Bertoloni (probable syntype), housed in MNHN. The illustration on the right was reproduced from Thomson (1856).

The two small holes on the longitudinal edges of the main type label suggest that it was a name label for the box the specimen was kept in, rather than a data label pinned to the specimen. It is likely that the name label was subsequently attached to the specimen when the latter was moved out of the original box. At the base of the label, Thompson wrote “♀ 200 Frs, ♂ £18”, indicating the existence of a male specimen of that species in the same box.

The male specimen referred to on this label is also housed in the MNHN collection, carrying a handwritten label with the price paid for the specimen of “£18”. This specimen is the same one described and illustrated by Westwood (1874) from the collection of James Aspinall Turner under the name of *Goliathus* (*Goliathinus*) *Fornassinii* (Fig. 5). The main data (*Fornassinii* / East Africa / ♂ Waller) and the perpendicular handwriting (“£18” and “Febry 11^th^ 1881 Sale”) on the specimen’s label are in different hands, the latter probably that of Thomson's. The information written to the perpendicular is consistent with the fact that the Turner collection was sold at auction on February 11^th^ 1881 at Stevens’ Auction Rooms, London (Horn *et al.*, 1990).

**Fig. 5.**
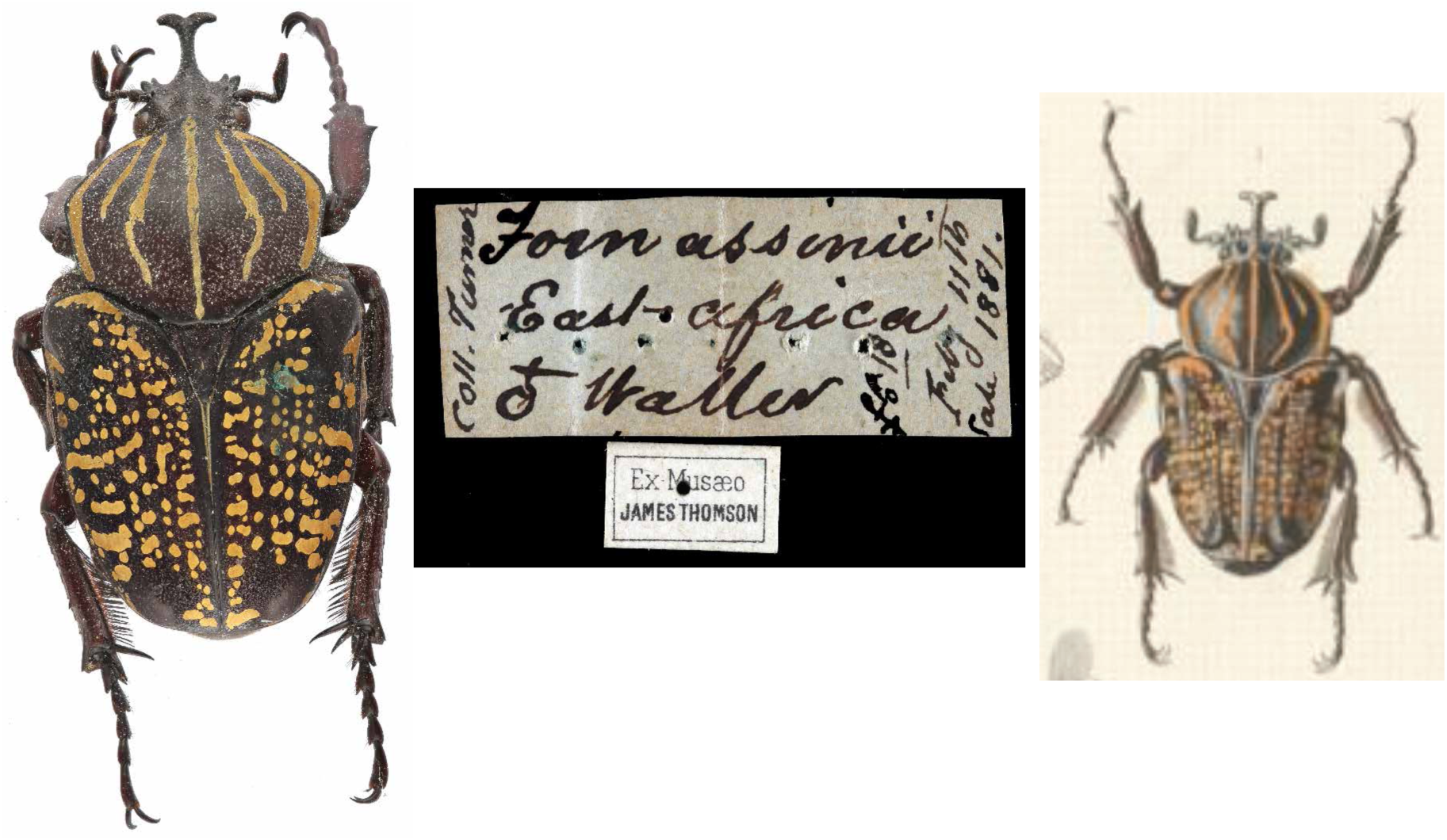
Westwood's specimen of *Fornasinius fornasini* Bertoloni, housed in MNHN. The illustration on the right was reproduced from Westwood (1874).

This male specimen was collected by Horace Waller in East Africa, according to the label data; or by John Kirk on the Zambezi River, according to Westwood (1874). Kirk was part of the David Livingstone Zambezi Expedition (1858–64), and many of his specimens would have been collected somewhere between Tete and Cahora Bassa in northern Mozambique (Livingstone, 1865). Waller was present in the same area at the same time (ca. 1861), as part of the Universities’ Mission to Central Africa, returning to England in 1862 (Rowley, 1866). Thus, it is unclear whether Kirk or Waller had collected Westwood’s specimen. Nevertheless, both the origin and structure of the cephalic horn of this male specimen (total length: 44 mm) are undoubtedly consistent with Betoloni’s *Fornasinius fornasini*.

Of note, Kraatz (1896) subsequently described *Fornasinius Westwoodi* based solely on Westwood’s illustration of *Goliathus* (*Goliathinus*) *fornassinii*, which he considered different from Bertoloni’s taxon (*“Ich nenne den Zambesi-Käfer vorläufig F. Westwoodi und mache auf die Form aufmerksam, welche sich leicht als eine eigene Art herausstellen könnte.”;* Kraatz, 1896) based on spurious and anecdotal characters (e.g., the small size, some details of the horn, and the extension of the pronotal bands). Thus, *Fornasinius westwoodi* Kraatz must be considered a synonym of *Fornasinius fornasini* Bertoloni.

Based on the aforementioned material and information, it is possible that Thomson assembled a box containing the two *Fornasinius fornasini* specimens, a female (syntype?) and a male, sometime before his death in 1897. This interpretation may explain why Thomson used the specific name *“Insignis”* for the box label containing the two specimens and complemented it with *“Fornassinii* Bert. Olim.”, to indicate the fact that the taxon was formerly known as *“Fornassinii”* at the time of his publication (Thomson, 1856).

A third male specimen has been illustrated by Nagai & Sakai (1998) under the name of *Fornasinius fornasini*. This specimen was collected at Lukuledi (IX.1911), a locality in the Mtwara region of southern Tanzania, at the border with Mozambique. This specimen is consistent with Bertoloni’s *Fornasinius fornasini*, i.e., in having an upturned, laterally compressed horn.

Finally, two further specimens (a male and a female; Fig. 6) are present in Gerhard Pross’ private collection, Germany. Both were collected in Ntandi, Mtwara Region, southern Tanzania, in XII.2012. Similar to Bertoloni’s male syntype, the male specimen in GPPC has a bulky body (total length: 55 mm) and a proportionally short frontal horn, decidedly bent upwards; prominently pointed ante-ocular frontal cornicles; and elongate, transversally truncated spines at the sides of the clypeal plate.

**Fig. 6.**
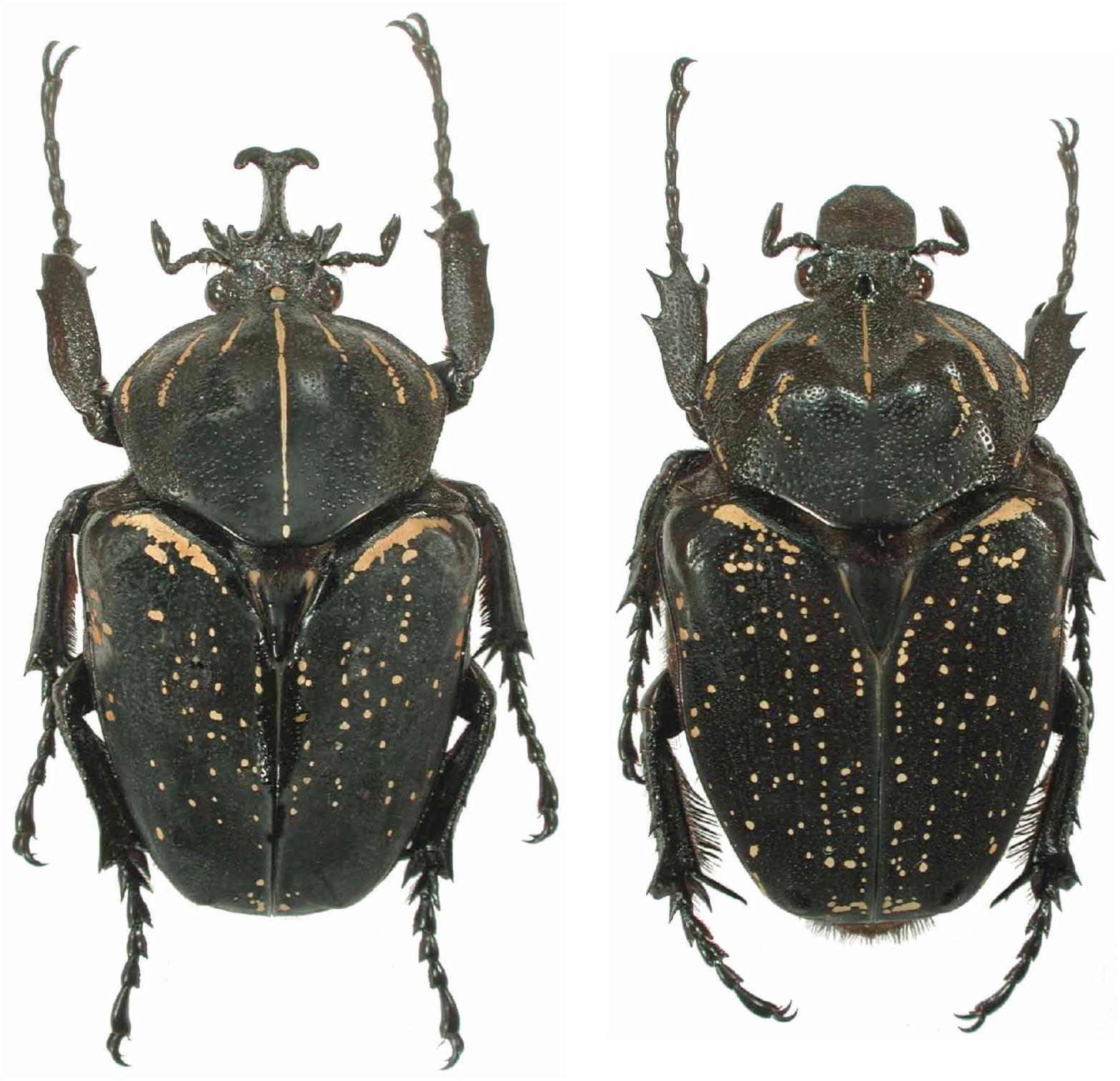
Male and female specimens of *Fornasinius fornasini* Bertoloni, housed in the private collection of Gerhard Pross, Germany.

The examination of the male specimen in Pross’ collection unequivocally confirmed the stability and peculiarity of the cephalic armature of *Fornasinius fornasini* Bertoloni (Fig. 7).

**Fig. 7.**
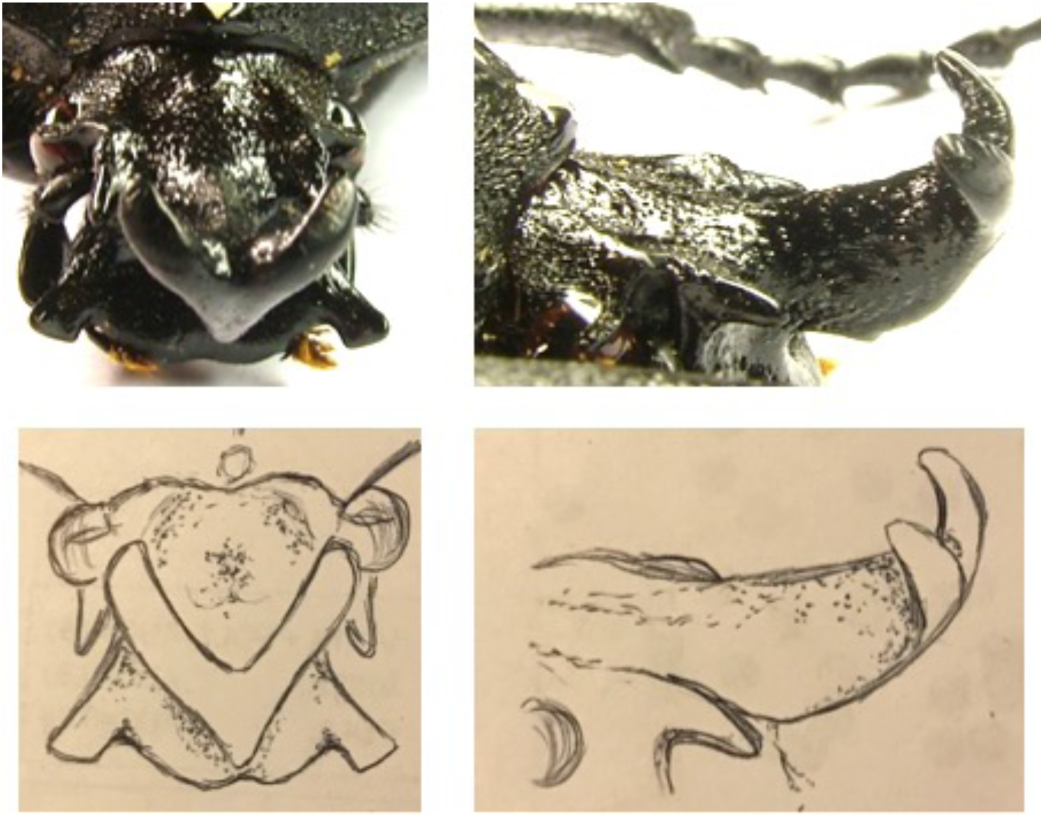
Head armature of a *Fornasinius fornasini* Bertoloni specimen, housed in the private collection of Gerhard Pross, Germany, seen from the front (left) or right side (right).

Based on the above, the following synonymy is established for Bertoloni’s taxon:

#### *Fornasinius fornasini* Bertoloni, 1853

= *Goliathus Fornasini* Bertoloni, 1853
= *Fornasinius insignis* Bertoloni, 1853
= *Goliathus Fornassinii* Thomson, 1856
= *Goliathus* (*Goliathinus*) *Fornassinii* Westwood, 1874
= *Fornasinius Westwoodi* Kraatz, 1896

#### Cephalic armature of the male

Frontal horn relatively short, compressed laterally, erected antero-dorsally, with terminal fork upturned and not bent downwards. Ante-ocular frontal cornicles prominent, pointed anteriorly. Clypeal plate with strongly indented, elongate, transversally truncated lateral spines.

#### Other characters

Pronotum with dense microsculpture, deeper and denser in females, with antescutellar area devoid of punctures. Male pro-tibia bidentate. Tarsi with short and robust tarsomeres. Pilosity black. Pronotum with tomentose, variably developed lateral margins and five discal bands. Tomentose marks orange.

#### Distribution

Mozambique (Inhambane, Tete), southern Tanzania (Mtwara).

### Identification of *Fornasinius fornasini auct*. (*nec* Bertoloni)

The lead author (M.D.P.) recently described a new *Fornasinius* species, *F. hermes* De Palma & Di Gennaro, 2017, from north-east Uganda. On that occasion, the taxa allied to *F. hermes* were illustrated without formally reviewing their status. Here, the taxa commonly referred to as *F. fornasini* (*sensu auct., nec* Bertoloni) are discussed.

In 1896, Kraatz described *Fornasinius Hauseri* based on two specimens, a male and a female, reportedly collected in the “Cameroon highlands” by Dr. Hauser (Kraatz, 1896). Unfortunately, the syntypes are neither in the Senckenberg Deutsches Entomologisches Institut (SDEI) of Müncheberg, Germany, where most of Kraatz’ types are housed, nor in Museum für Naturkunde of Berlin (MNHB), Germany, where most of Hauser’s material is housed. Nevertheless, Kraatz’ description and illustration of *Fornasinius hauseri* are sufficiently detailed to enable its identification.

The male syntype of *Fornasinius hauseri* measures 42 mm (Kraatz, 1896). According to the description and illustration (Fig. 8), the cephalic horn has a relatively broad base and a tapering, carinate distal segment; this structure gives the horn a triangular outline when observed from above. Notably, the horn projects dorsally from the frons, but bends downwards for most of its length. The features of the cephalic horn of *Fornasinius hauseri* are, therefore, markedly different from those of *Fornasinius fornasini* Bertoloni, whereas they are fully consistent with those of *Fornasinius fornasini auct*. (*nec* Bertoloni).

**Fig. 8.**
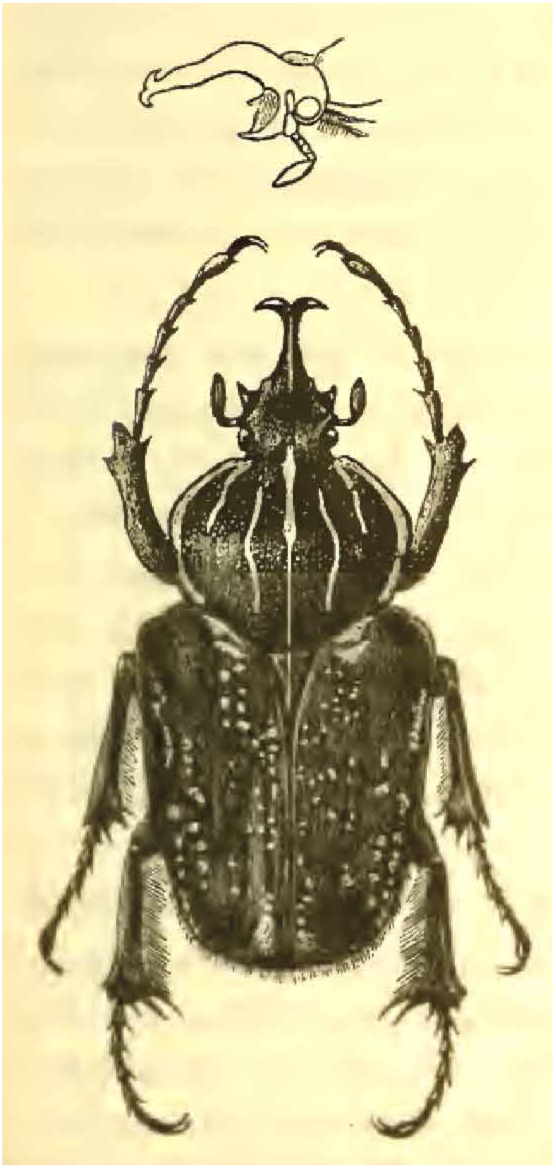
Original drawing of *Fornasinius hauseri* Kraatz. The illustration was reproduced from Kraatz (1896).

Kraatz (1896) compared *Fornasinius hauseri* with *Goliathinus aureosparsus* (Neervoort Van de Poll, 1890). He distinguished the two based on the punctuation of the pronotum, which is deeper in *hauseri* (*“aufserdem ist der Thorax von aureosparsus auf der Mitte glatt und nur zerstreut punktirt, während der Thorax der Kamerun-Art in der Mitte ziemlich dicht punktirt ist.”*] Kraatz, 1896) and the colour of the pygidial pilosity, which is reddish in *aureosparsus* (*“mit fuchsrother Hinterleibsspitze”;* Kraatz, 1896), among other characters.

Four years after Kraatz’ paper, Heath (1900a) described *Goliathinus* (*Sphyrorrhina*) *Wisei*, based on a single male specimen (holotype; Fig. 9), currently housed in the Natural History Museum of London (BMNH), UK. The specimen was collected in Nengia (= Kitui region), Kenya. Immediately after the description of *Goliathinus* (*Sphyrorrhina*) *wisei*, Heath published a note in which he acknowledged the similarity of his taxon with *Fornasinius hauseri* (Heath, 1900b), but insisted on a number of distinguishing characters that at best amount to intraspecific variation. Of note, the holotype of *Goliathinus X* (*Sphyrorrhina*) *wisei* is a relatively large specimen (53 mm) compared to the male syntype of *Fornasinius hauseri*. Although Kraatz’ syntypes, could not be traced, the available information suggests that *Goliathinus* (*Sphyrorrhina*) *wisei* is a junior synonym of *Fornasinius hauseri*, which is the first available name for *Fornasinius fornasini auct*. (*nec* Bertoloni).

**Fig. 9.**
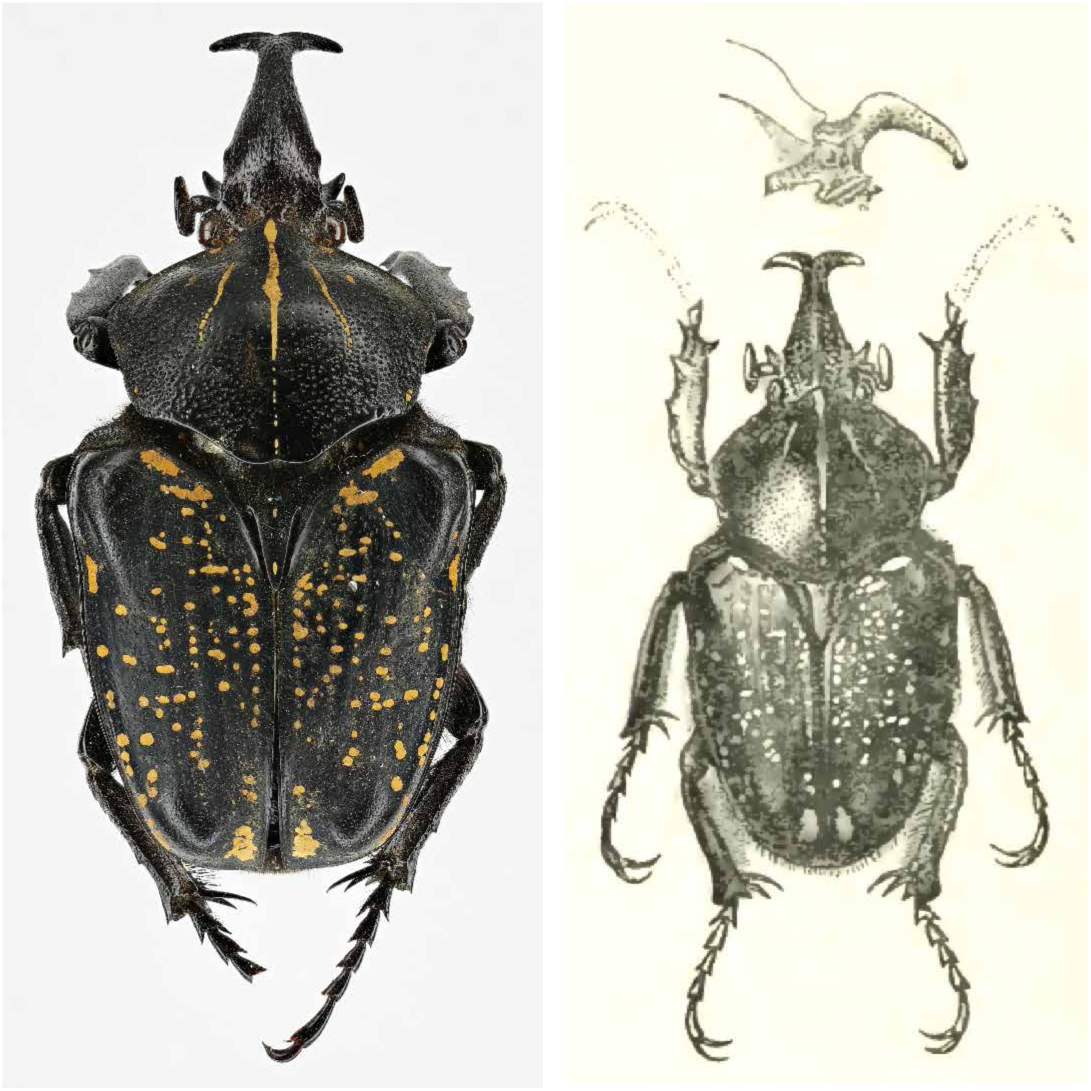
*Goliathinus* (*Sphyrorrhina*) *wisei* holotype (left), housed in BMNH (NHMUK010368075), and original drawing (right), reproduced from Heath (1900a).

Numerous specimens have been collected broadly in south-central Kenya that conform to the description of Kraatz’ and Heath’s taxa. Conversely, there are no reliable records from Cameroon, with the possible exception of one specimen similar to Kraatz’ syntype, which has been illustrated by Beinhundner (2017) under *F. fornasini var. wisei* (specimen n. 8). Although the very disjunct distribution of *Fornasinius hauseri* casts doubts on the Cameroonian origin of Kraatz’s syntypes, the possibility that a rare population of this species exists (or had existed) in Cameroon cannot be excluded. For example, *Goliathus goliatus* (Drury, 1770) is found in the mountainous regions of both Cameroon and Kenya, with little intra-specific variability.

Based on the above, the following synonymy is established for Kraatz’ taxon:

#### *Fornasinius hauseri* Kraatz, 1896, *bon. sp*

*= Goliathinus* (*Sphyrorrhina*) *Wisei* Heath, 1900
= *Fornasinius fornasini auct*. (*nec Bertoloni*)

#### Cephalic armature of the male

Frontal horn with a broad base projecting dorsally from the frons and a tapering, carinate distal segment projecting downwards. In rare cases, the cephalic horn may display lateral spines, although less developed than in *F. aureosparsus* and *F. hermes*. Ante-ocular frontal cornicles absent in most specimens, or poorly developed. Clypeal plate with softly indented, elongate lateral spines.

#### Other characters

Microsculpture, legs and pilosity as in *F. fornasini*. Pronotum with tomentose, variably developed lateral margins and five discal bands. Tomentose areas dark orange, generally sparse and reduced on elytra.

#### Distribution

south-central Kenya (Kitui and neighbouring regions), Cameroon?

### *Fornasinius hauseri* populations of Lake Victoria and neighboring regions

A well-differentiated population of *Fornasinius hauseri* is distributed in the surroundings of Lake Victoria, stretching eastward to northern Tanzania (De Palma & Di Gennaro, 2017). This taxon was described by Preiss (1904) from Süd-Nyansa (Rwanda) under the name of *Fornasinius Hirthi* (Fig. 10).

**Fig. 10.**
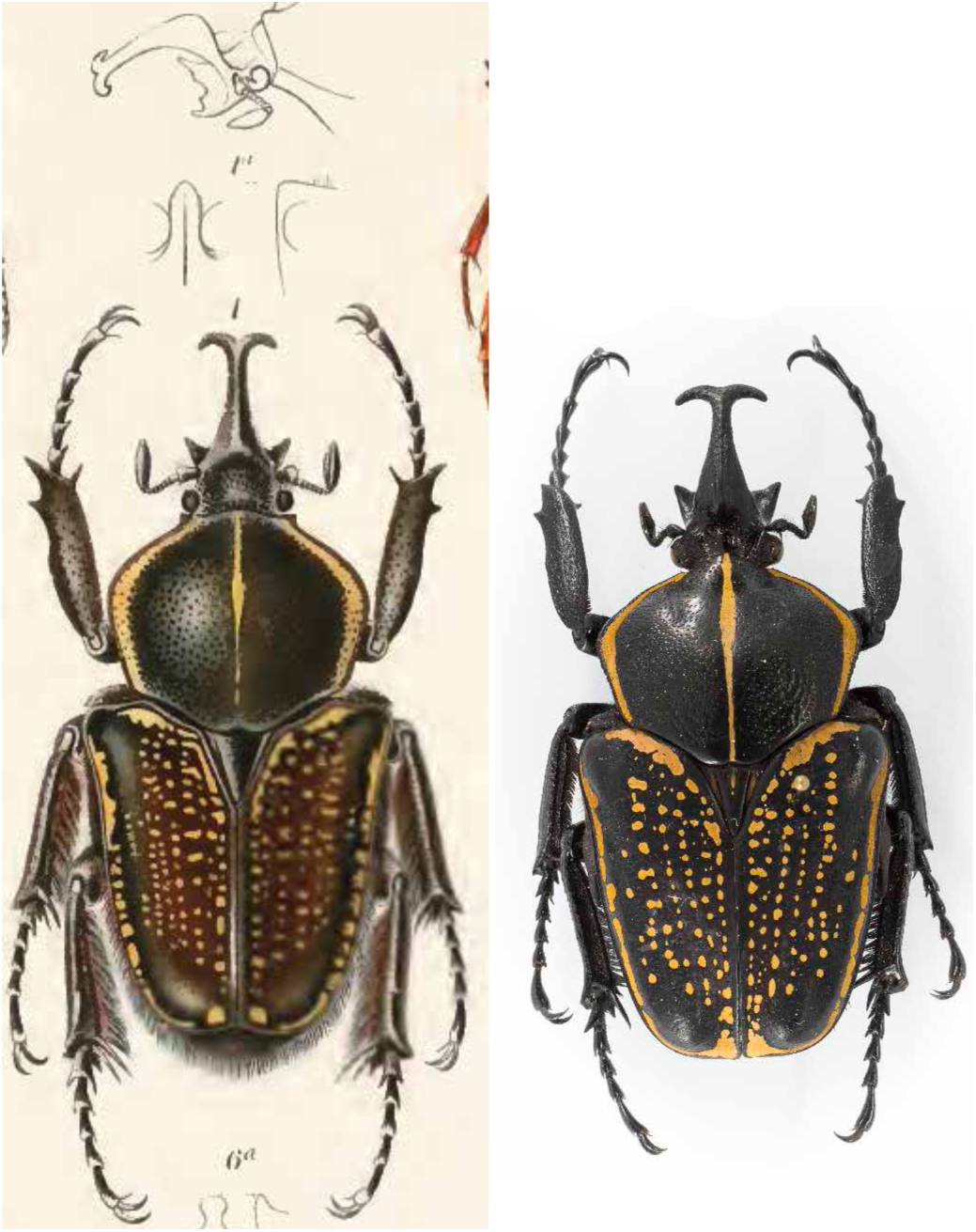
Original drawing of the *Fornasinius hirthi* syntype (left), reproduced from Preiss (1904). A specimen from North-East Tanzania is shown on the right.

Although the external characters (e.g., structure of the cephalic horn and denticulation of the tibia) are similar to *Fornasinius hauseri, Fornasinius hirthi* displays a stable coloration throughout its broad distributional range. *Fornasinius hirthi* is, therefore, raised from the synonymy with *Fornasinius fornasinius* and ranked subspecifically under *Fornasinius hauseri*. To this day, this is the most collected and best-known *Fornasinius* taxon.

Kolbe (1913; 1914) described five taxa (as “varieties”) under *Fornasinius insignis* (*nec* Bertoloni): *F. paradoxus, F. pauxillus, F. mixtus, F. transitivus*, and *F. infradentatus*. The series was collected in Lake Victoria (Ukerewe Is.) and north-east Tanzania (Usambara Mts.); most of the types are housed in MNHB. Upon examination, it has emerged that they all belong to *Fornasinius hauseri* ssp. *hirthi*. Kolbe’s varieties were described from specimens of different size and based on color variation as well as the structure of the clypeal horn, which varies allometrically in this species (Weibes, 1968).

The following synonymy is, therefore, established for Preiss’ taxon:

#### *Fornasinius hauseri* ssp. *hirthi* Preiss, 1904, *stat. rev*

= *Fornasinius Hirthi* Preiss, 1904
= *Fornasinius insignis* var. *paradoxus* Kolbe, 1913
= *Fornasinius insignis* var. *mixtus* Kolbe, 1913
= *Fornasinius insignis* var. *pauxillus* Kolbe, 1913
= *Fornasinius insignis* var. *transitivus* Kolbe, 1913
= *Fornasinius insignis* var. *infradentatus* Kolbe, 1913
= *Fornasinius fornasini auct*. (*nec Bertoloni*)

#### Cephalic armature of the male

As in the nominate form.

#### Other characters

Microsculpture, legs and pilosity as in the nominate form. Pronotum with thick, tomentose lateral margins and a thick, longitudinal central band. Complete or partially complete line of tomentose marks on the elytral margins. Elytral maculation generally more dense than in the nominate form. Tomentose marks orange.

#### Distribution

Rwanda, Burundi, western Kenya, southern Uganda, north-east Democratic Republic of the Congo, northern Tanzania.

A third subspecies is to be described from south-east Democratic Republic of the Congo.

### Note

The taxonomic changes proposed in this article, as well as the description of new taxa in the genus *Fornasinius*, will be published in a forthcoming article to appear in *Entomologia Africana* (ISSN: 1371–7057) in 2018.

## Methods

The original spelling (e.g., capitalization of the specific name) is given for each taxon at the first occurrence in the text.

The length of the specimens is given from the tip of the clypeal horn to the tip of elytra.

This taxonomic study has been undertaken by reviewing all the relevant literature and by examining primary types and other specimens deposited in the following public collections:

BMNH: Natural History Museum, London, UK.
MNHB: Museum für Naturkunde, Berlin, Germany.
MNHN: Muséum national d'Histoire naturelle, Paris, France.
SDEI: Senckenberg Deutsches Entomologisches Institut, Müncheberg, Germany.
UNIBO: Dipartimento di Scienze Biologiche, Geologiche e Ambientali, Universitá di Bologna, Italy.

Additional specimens were studied from the private collections of M.D.P (Switzerland) and Mr. Gerhard Pross (Germany).

## Acknowledgements

M.D.P. is indebted to Didier Camiade and Gerhard Beinhundner for the insightful discussions on the identity of *Fornasinius fornasini* Bertoloni. Prof. Mario Marini is warmly thanked for providing photos of the *Fornasinius fornasini* syntypes housed in UNIBO; Antoine Mantilleri and Jean-Philippe Legrand for providing photos of specimens housed in MNHN; Bernd Jaeger, Bernhard Schurian and Joachim Willers for providing information on, and photos of, specimens housed in MNHB; Kostiantyn Nadein for searching for the *F. hauseri* syntypes in the SDEI collections; and Gerhard Pross (Germany) for providing photos of specimens in his collection.

